# Neural Dynamics Underlying Successful Auditory Short-Term Memory Performance

**DOI:** 10.1101/2023.06.30.547170

**Authors:** Ulrich Pomper, Lorenza Zaira Curetti, Maria Chait

## Abstract

Listeners often operate in complex acoustic environments, consisting of many concurrent sounds. Accurately encoding and maintaining such auditory objects in short-term memory is crucial for communication and scene analysis. Yet, the neural underpinnings of successful auditory short-term memory (ASTM) performance are currently not well understood.

To elucidate this issue, we presented a novel, challenging auditory delayed match-to-sample task while recording MEG. Human participants listened to “scenes” comprising 3 concurrent tone pip streams. The task was to indicate, after a delay, whether a probe stream was present in the just-heard scene. We present three key findings: First, behavioural performance revealed faster responses in correct vs. incorrect trials as well as in ‘probe present’ vs ‘probe absent’ trials, consistent with ASTM search. Second, successful compared to unsuccessful ASTM performance was associated with a significant enhancement of event-related fields and oscillatory activity in the theta, alpha, and beta frequency ranges. This extends previous findings of an overall increase of persistent activity during short-term memory performance. Third, using distributed source modelling, we found these effects to be confined mostly to sensory areas during encoding, presumably related to ASTM contents per-se. Parietal and frontal sources then became relevant during the maintenance stage, indicating that effective STM operation also relies on ongoing inhibitory processes suppressing task irrelevant information.

In summary, our results deliver a detailed account of the neural patterns that differentiate successful from unsuccessful ASTM performance in the context of a complex, multi-object auditory scene.

## 1. Introduction

Auditory short-term memory (ASTM) refers to the ability to maintain auditory information over durations of a few seconds and is an important requirement for the analysis of acoustic scenes (Wilsch and Obleser, 2016). As such, it is involved in basic, yet vital tasks such as detecting potentially threatening changes in our acoustic environment, as well as highly complex functions like verbal communication or musical performance.

Recent years have shown an increasing interest in human ASTM (Arnott et al, 2005; Buchsbaum et al., 2005; Cappotto et al., 2021; Kumar et al., 2016; Lim et al, 2022; Noyce et al., 2022; Papagno et al., 2017), particularly in the context of electrophysiological oscillatory correlates (Kaiser, 2015; Kumar et al., 2021; Weisz et al., 2020; Wilsch and Obleser, 2016). Whilst previous work has predominantly focused on contrasting low- and high-load ASTM tasks, here we ask what patterns of brain activity distinguish successful vs. unsuccessful ASTM performance whilst the task, and stimuli presented to participants, remain fixed.

Accumulating work has identified core components of ASTM, primarily comprising auditory cortex as well as regions in the prefrontal (Albouy et al., 2017; Huang et al., 2013; Kumar et al., 2021, 2016) and parietal cortices (Koelsch et al., 2009; Leung et al. 2011). It is suggested that while the contents of ASTM are represented in sensory areas, prefrontal regions are performing executive control functions coordinating the encoding, maintenance and retrieval of stored information (Spitzer and Blankenburg, 2012; Sreenivasan et al., 2014; Deutsch et al., 2023).

Functionally, a common observation is of increases in event-related EEG and MEG activity during high compared to low ASTM load maintenance (Grimault et al., 2014; Huang et al., 2016; Nolden et al., 2013), which is suggested to originate from elevated firing of neurons in memory relevant areas (Huang et al. 2016).

Similar to observations in the context of visual short-term memory (Ratcliffe et al., 2022; Roux and Uhlhaas, 2014), oscillatory neural activity (as measured with EEG and MEG) is suggested to play an important role in encoding and maintaining information in ASTM (Kaiser, 2015; Wilsch and Obleser, 2016; Weisz et al., 2020; Kumar et al., 2021; Figure 1). In particular, increased theta power (4-7 Hz) has been observed in sensory areas during encoding of ASTM, suggested to reflect chunking of incoming information (Teng et al., 2017), as well as in parietal cortex during maintenance, indicative of ASTM manipulation (Albouy et al., 2017, 2022). Likewise, increase in alpha-band activity (8-13 Hz) over posterior areas has been repeatedly associated with ASTM tasks, interpreted to reflect the suppression of task-irrelevant, potentially distracting information (Obleser et al., 2012; Wisniewski et al., 2017; Wöstmann et al., 2017). Further, enhancements of beta-band power (14-30 Hz) in temporal regions have been observed with increasing levels of ASTM load (Leiberg et al., 2006), and associated with binding of auditory short-term memory contents in sensory areas (Weiss and Mueller, 2012).

**Figure 1.**
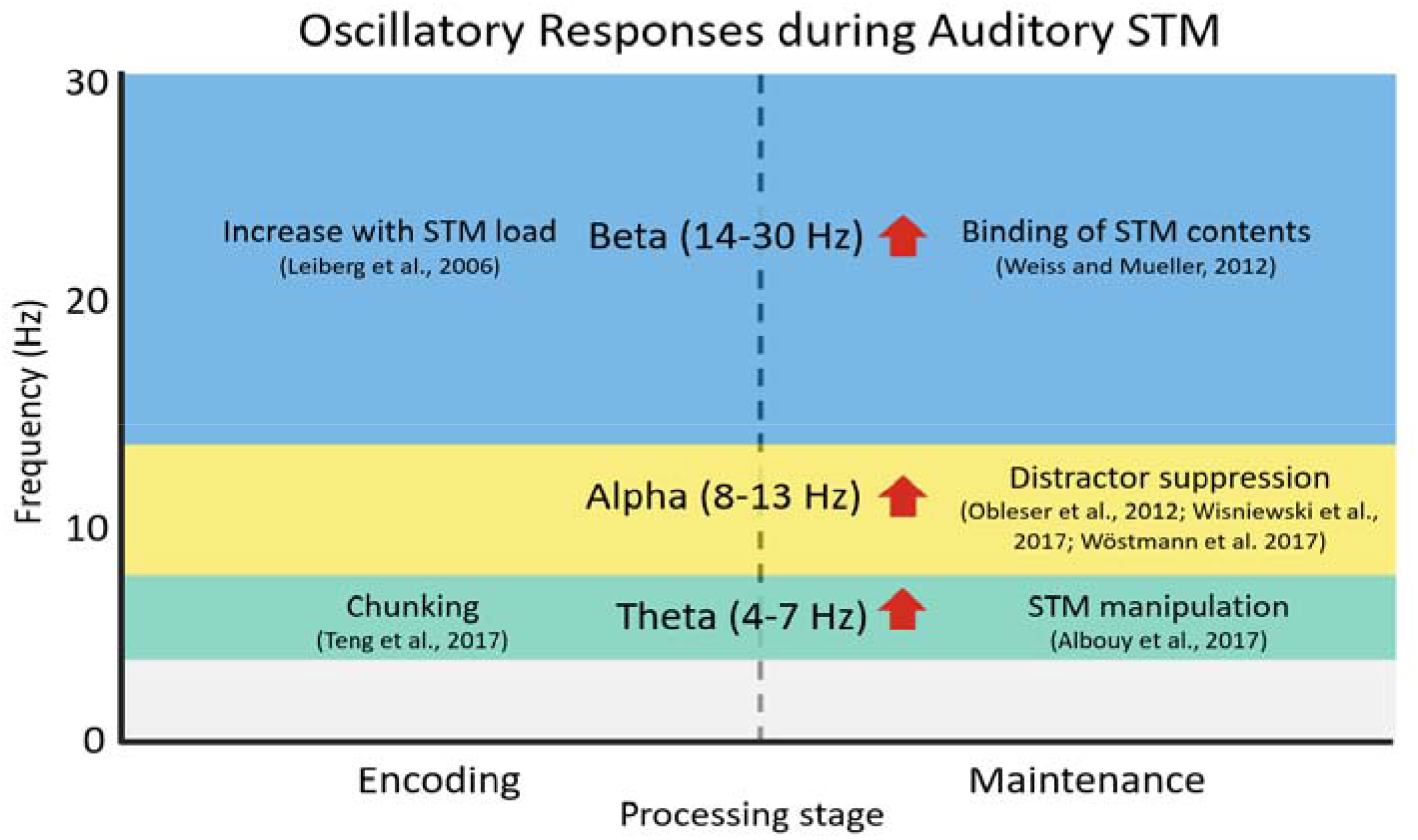
Overview of common oscillatory responses during auditory STM tasks reported in previous studies.

Overall, these findings suggest a rich spatio-spectral activity pattern associated with ATSM, in which different functional aspects of the task are carried out by localized activity in specific frequency bands.

However, while recent studies have investigated how these oscillatory signatures are affected by task requirements (e.g. Albouy et al., 2017; Dimakopoulos et al, 2022; Wöstmann et al. 2017) or STM load (Leiberg et al., 2016; Obleser et al. 2012; Kosachenko et al., 2023), it is currently unknown how they might relate to successful versus unsuccessful ASTM performance where task load per se is constant across trials (for examples in the visual domain, see e.g. Lenartowicz et al., 2021; Mapelli & Ozkurt, 2019; Proskovec et al., 2018;).

Here, using MEG, we examined brain activity associated with (ultimately) correct vs. incorrect responses in a challenging short-term memory task. Stimuli (Figure 2) were artificial ‘acoustic scenes’ consisting of three concurrent auditory ‘objects’, akin to a mix of auditory sources in everyday listening environments. Listeners were required to memorize the sounds, and, following a silent retention interval, determine whether a probe sound was present in the just-heard scene. We explored the evoked and oscillatory activity during encoding and maintenance, identifying a distributed pattern of neural dynamics, which differentiates subsequently successful versus failed memory performance.

**Figure 2.**
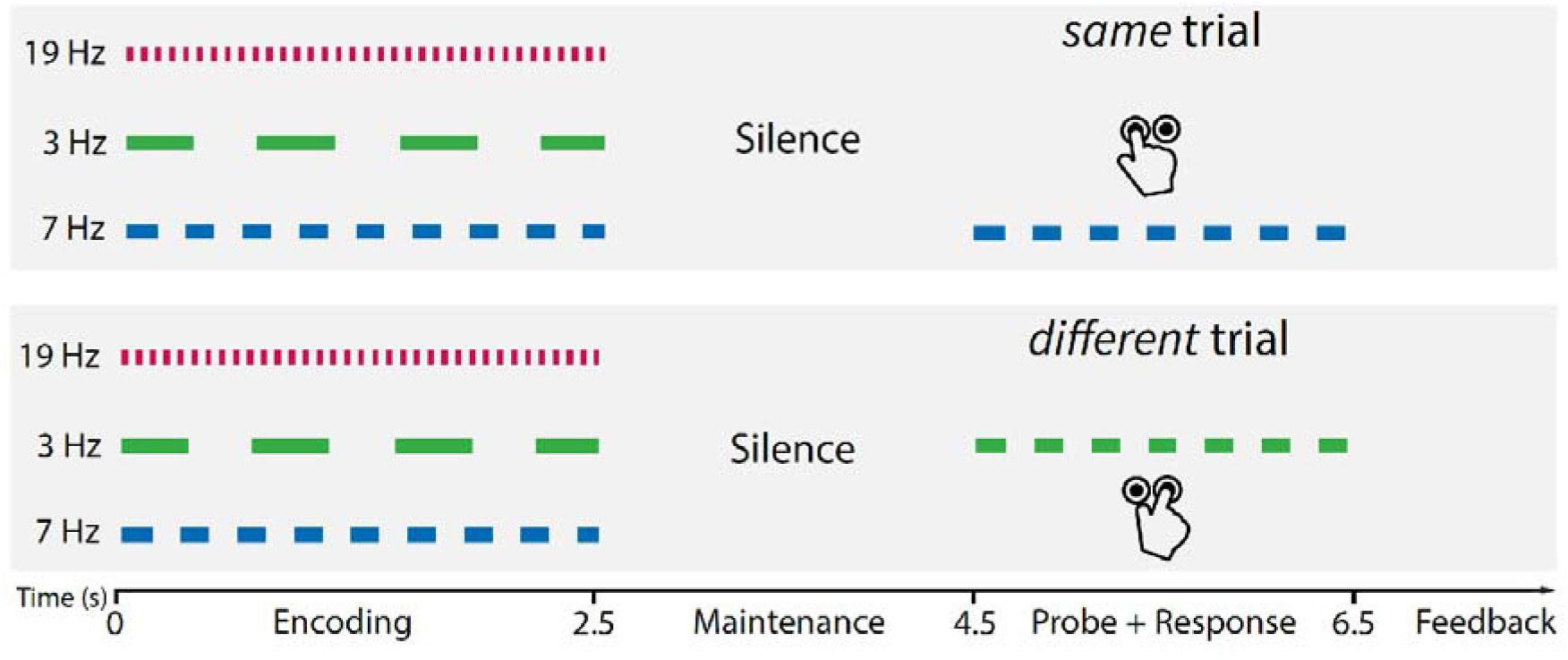
Schematic trial structure. Each trial started with an ‘encoding’ stage lasting 2500 ms, during which an auditory scene consisting of 3 concurrent streams was presented. Stream rates were chosen from a set of 3 values: 3 Hz, 7 Hz, and 19 Hz. Carrier frequencies (indicated by the red, green and blue colours) were randomly assigned to each stream on a trial-by-trial basis (taken from a pool of 11 values). The Encoding stage was followed by a 2000 ms silent Maintenance stage. After this, a probe stimulus was presented for 2000 ms. This probe was either identical to one of the streams presented during encoding (*same* trial; top panel), or a combination of one of the assigned carrier frequencies with one of the two AM rates with which it was not originally paired (*different* trial; bottom panel). For instance, in the example plotted here the green carrier frequency, originally presented (in the encoding stage) at 3 Hz, is now paired with the 7 Hz rate. Participants were asked to provide a speeded response, indicating whether the probe stimulus was *same* or *different*, using one of two buttons.

## 2. Methods

### 2.1. Participants

Twenty-eight right handed (Edinburgh Handedness Inventory; Oldfield, 1971), paid participants took part in the study. All reported no neurological illness or hearing deficits. One participant was excluded due to not being able to properly perform the task, despite repeated training. Four other participants were excluded due to exceptionally poor behavioural performance (mean response times of > 2000 ms compared to 1007 ms for the rest of the participants). Therefore, data from twenty-three participants (16 females, mean age 23.6 years) are presented. The study was conducted in accordance with the research ethics committee of the University College London and all participants provided written informed consent.

### 2.2. MEG recordings

MEG was recorded continuously (600 Hz sampling rate; 100 Hz hardware low-pass filter) using a CTF-275 MEG system, with 274 functioning axial gradiometers arranged in a helmet shaped array. Electrical coils were attached to three anatomical fiducial points (nasion and left and right pre-auricular), to continuously monitor the position of each participant’s head with respect to the MEG sensors. During the recording, participants were seated in an upright position in a magnetically shielded room. Sounds were presented binaurally via insert earphones (E-A-RTONE 3A 10, Etymotic Research), with sound intensity adjusted individually to a comfortable level. To minimize eye movements, a black fixation cross on a grey background was presented throughout the experiment via a screen located about 60 cm in from of the participant’s eyes.

### 2.3. Task and Stimuli

At the beginning of each trial, participants were presented with a 2500 ms long auditory ‘scene’ consisting of three simultaneous streams of repetitive pure tones (Figure 2; ‘encoding’ stage). Each stream had a unique frequency and temporal rate (TR). Rates were 3 Hz, 7 Hz and 19 Hz, defined by the duration of a single tone and the following inter-tone interval (each 166.5 ms, 71.45 ms, and 26.15 ms respectively, i.e. SOAs of 333 ms, 142.9 ms and 52.3 ms, respectively). Each tone had a rise and fall time of 5 ms. Frequencies were randomly chosen from a pool of eleven logarithmically spaced values (200, 292, 405.1, 542, 708.4, 910.7, 1157, 1456, 1820, 2262, and 2799 Hz). As a consequence, the rate content of the scene was fixed but, on each trial,, these were paired with different carrier frequencies. The ‘encoding’ stage was followed by a silent interval of 2000 ms duration, during which participants had to maintain the auditory scene in memory (‘maintenance’ stage). After the gap, a single auditory stream was presented as a probe for 2000 ms. In 50% of trials, this probe was identical to one of the three initially presented streams (*same* trial), and in 50% of trials it consisted of one of the three presented carrier frequencies combined with one of the presented rates, that it was not previously paired with (*different* trial). Each participant completed 336 trials in total (56 per probe rate for both the *same* and *different* conditions).

Participants were instructed to provide a speeded response via one of two buttons with their right hand, to indicate whether the probe belonged to the *same* or *different* category. After every button press, visual feedback (lasting 300 ms) was given on whether the answer was correct or incorrect. Following the offset of the feedback was an inter-trial interval varying uniformly between 700 and 2000 ms (in steps of 100 ms,).

The fact that the probe consisted of a previously presented frequency and rate rendered it always partially ‘familiar’, which increased task difficulty. To perform correctly, participants had to accurately bind the appropriate frequencies and rates during the ‘encoding’ stage and maintain this information in memory until probed. Pilot experiments revealed that performance accuracy is on average at around 70%, suggesting it is a sufficiently challenging task to probe short-term memory and failures thereof (see also Figure 3B).

**Figure 3.**
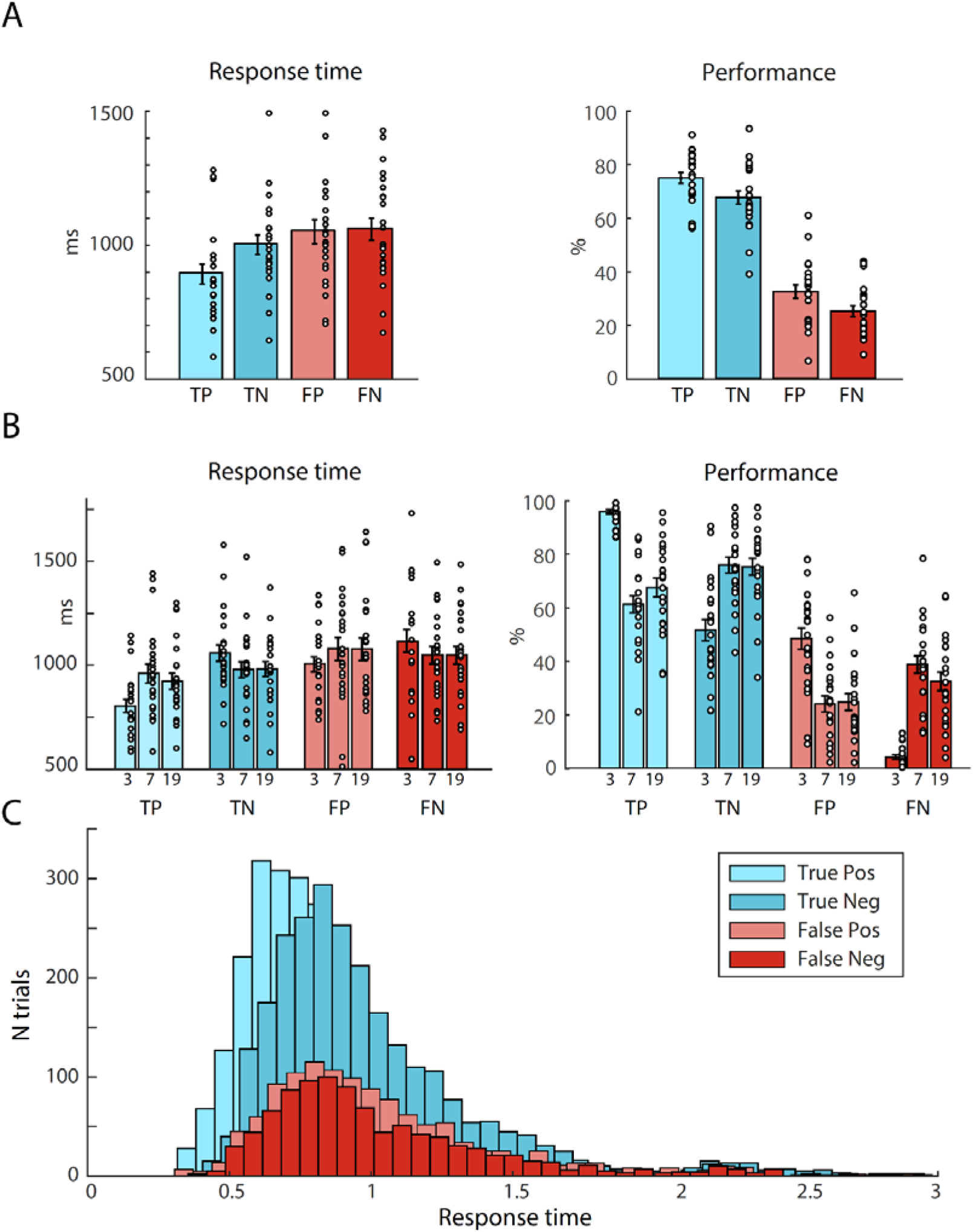
Behavioural results. **(A)** Data pooled across probe-rates, separately for trials with true positive (TP), true negative (TN), false positive (FP), and false negative (FN) outcomes Left: Average response times. Right: Performance (% of *same* [TP + FN] and *different* [TN + FP] trials). Circles indicate individual participants, error bars represent SEM. **(B)** Same as A, with separate bars for trials with probes of 3 Hz, 7 Hz and 19 Hz AM **(C)** Probability density function of RTs separately for TP, TN, FP, and FN responses.

### 2.4. Behavioural analysis

Prior to the statistical analysis, outlier trials with RTs deviating more than 2.5 SDs from the mean were excluded per participant and condition. Mean percentage of true-positives (TP), true-negatives (TN), false-positives (FP), and false-negatives (FN), as well as response times (RTs) were computed separately for each condition. For statistical analysis, performance and RT data were each subjected to a repeated-measures analysis of variance (RMANOVA), including the within-participant variables outcome (TP, TN, FP, FN) and rate (3, 7, 19 Hz).

### 2.5. MEG analysis

#### Preprocessing

Off-line, MEG was processed using Matlab (The MathWorks Inc., Natick, MA, USA), and FieldTrip (Oostenveld et al., 2011). First, data were epoched from -1000 ms to 6000 ms around the onset of auditory scenes. Thus, each epoch consists of a 1000 ms ‘prestimulus’ stage, a 2500 ms ‘encoding’ stage, a 2000 ms ‘maintenance’ stage, and a 2000 ms ‘probe’ stage. Trials with power that deviated from the mean by more than twice the standard deviation (across all sensors and time points) were deemed outliers and discarded from further analyses. Artifacts such as slow drifts and 50 Hz line noise were removed using DSS (denoising source separation; de Cheveigne and Parra, 2014), and epochs were baseline corrected relative to a 500 ms prestimulus interval.

All data were then separated into *correct* (TP, TN) and *incorrect* (FP, FN) conditions, based on the behavioural response. Within each participant, the number of *correct* trials was larger than that of *incorrect* trials. As the signal-to-noise ratio increases with the number of trials used in averaging procedures such as the presently investigated event-related fields and event-related oscillations, this difference in the number of trials between conditions potentially confounds their comparison. To ensure similar signal-to noise ratio in both conditions, we selected a subset of *correct* trials for each participant, equal in number to the *incorrect* trials. Since outlier trials (+/- 2.5 SD away from the mean) at both tails of the RT distribution were already discarded beforehand, RTs can be used as a marker for rating confidence, as well as to disentangle true *correct* responses from guesses (Ratcliff, 1993; ‘true’ *correct* responses are expected to be faster than ‘guess’ responses). Thus, we equated the numbers of trials by ordering all *correct* trials according to response time and discarding the surplus trials from the slow tail of the distribution.

#### Time domain analysis

First, we investigated potential differences between event related fields (ERFs) during *correct* versus *incorrect* trials. As a measure of global instantaneous power, for each participant, the root mean square (RMS) amplitude across all channels was calculated separately for the *correct* and *incorrect* trials. To identify time windows where *correct* trials differed from *incorrect* trials, a bootstrapping procedure (10000 iterations) was performed on the difference in RMS time-course between the two conditions. On each iteration, the data were resampled by choosing random sets of 23 participants (with replacement). For each timepoint, we computed the mean response across participants, and any time-point yielding above 95% of iterations in the same direction (positive or negative) was deemed significant (Figure 4A). This analysis yielded widespread significant differences throughout the encoding and maintenance stages (Figure 4A). To identify the channels that contribute to this global effect, a cluster-based permutation procedure (Maris and Oostenveld, 2007; 1000 permutations, min. = 3 channels) was used. The ERF cluster correction was performed on the spatial (i.e. channels) dimension only, separately for the encoding and maintenance stages. Finally, cortical sources for any observed differences were estimated via the multiple-sparse priors (MSP) method implemented in SPM12. Participant-specific forward models were computed using a Single Shell model and sensor positions projected onto an MNI space template brain by minimizing the sum of squared differences between the digitized fiducials and the MNI template. Data for the encoding and maintenance stages (i.e. from 0 to 4500 ms following stimulus onset) were inverted conjointly and pooled across *correct* and *incorrect* trials. Grand average source solutions were constrained by the restriction that included sources have to be present in each participant, which has been shown to improve group-level statistical power (Litvak and Friston, 2008). After inversion, source estimates were averaged over intervals of significant differences between *correct* and *incorrect* trials observed via the RMS analysis, projected to a 3D source space (MNI space, constrained to gray matter), and smoothed (8-mm full width at half maximum (FWHM) Gaussian smoothing kernel) to create images of source activity for each participant. Given that the goal of source reconstruction was to localize the neural generators of sensor-space effects previously identified as significant, statistical maps of source activity are displayed with uncorrected voxel-wise thresholds (Gross et al., 2013).

**Figure 4.**
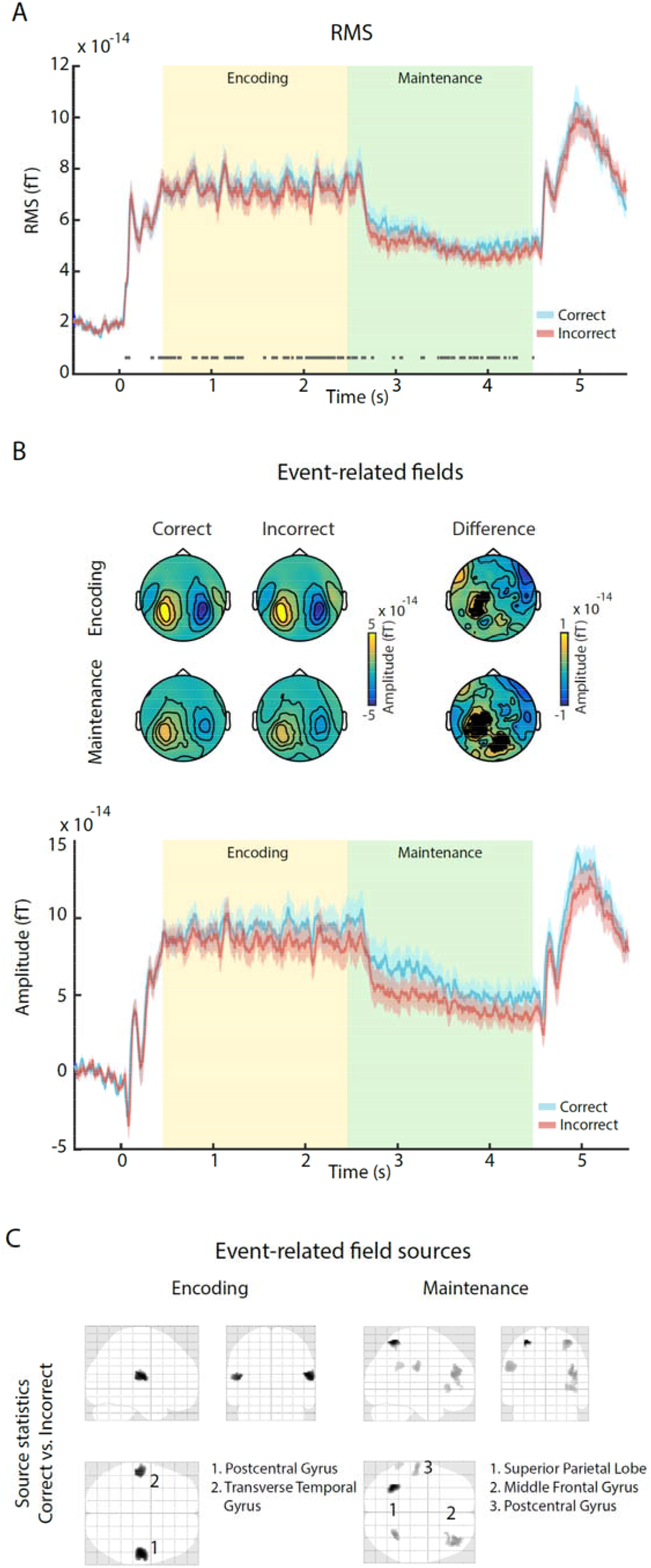
Evoked responses across the entire experimental trial. **(A)** Global instantaneous power (root-mean square activity across all channels), for *correct* (blue) and *incorrect* (red) trials. The horizontal grey line indicates stages of significant differences as observed via a bootstrap permutation analysis. Yellow and green shaded areas indicate Encoding and Maintenance stages, respectively. **(B)** Top: Scalp topographies for *correct* (left) and *incorrect* (middle) trials, as well as their difference (right). Black dots indicate channels exhibiting significant differences. Bottom: Event-related field traces for *correct* (blue) and *incorrect* (red) trials (mean over channels on the left). Yellow and green shaded areas indicate Encoding and Maintenance stages, respectively. **(C)** Source-level difference between *correct* and *incorrect* trials, separately for the encoding (left) and maintenance stage (right). Numbers indicate significant clusters (p<0.05), sorted by p-values from low to high. Insets indicate anatomical areas containing the peak voxel of each cluster.

#### Analysis of oscillatory power

To study differences in oscillatory neural activity between *correct* and *incorrect* trials, all data were transformed into time–frequency domain by applying a sliding window Fourier transform with a single Hanning taper. Power at frequencies from 1 to 30 Hz was computed in 0.5 Hz steps, using a fixed time window (400 ms) and a fixed frequency smoothing (1 Hz). For baseline correction, oscillatory power in each trial was normalized relative to the interval from -400 to -200 ms (indicating the centre of the sliding time window, thus including data from -600 to 0 ms prior to the beginning of the trial) before the onset of the auditory stimulus.

For statistical comparison between *correct* and *incorrect* trials, cluster based permutation procedures were employed (Maris and Oostenveld, 2007; 1000 permutations, min. = 3 channels), with cluster-correction in the temporal and spatial (i.e. channel) domain. Statistics were calculated separately for the ‘prestimulus’, ‘encoding’ and ‘maintenance’ stages, within the delta (1-4 Hz), theta (4-7 Hz), alpha (8-13 Hz), and low beta (14-20 Hz) range.

Similar to the analysis of ERF responses, we estimated the sources underlying the effects in oscillatory power and phase observed at the sensor-level. Here, we used the minimum-norm method (Hämäläinen and Ilmoniemi, 1994) implemented in SPM 12, which has been shown to be suitable for the localization of induced oscillatory activity (Halder et al., 2019). Data for the encoding stage (from 500 to 2500 ms, i.e. excluding the evoked response from stimulus onset) and maintenance stage (from 2500 to 4500 ms) were first inverted separately, and independently for the theta (4-7 Hz), alpha (8-14 Hz), and lower beta (15-20 Hz) frequency ranges. Source solutions were pooled across *correct* and *incorrect* trials and constrained to be consistent across participants. Significant effects from sensor-space were localized within the brain (in MNI space, constrained to grey matter) after summarizing source power in the specific time-frequency range of interest for each participant using a Morlet wavelet projector (Friston et al., 2006).

#### Power-spectral density analysis

In addition to effects on ongoing endogenous neural oscillations, the presentation of regular auditory inputs has been shown to cause entrainment of neural activity (Falk et al., 2017, Kösem et al., 2018, van Bree et al., 2020). As an indication of successful versus unsuccessful ASTM performance, we speculated that the strength of entrainment during encoding reflects the quality of sensory processing, while its continuation during active maintenance might correlate with the fidelity of ASTM representations and subsequent performance. As a measure of the fidelity of encoding and maintenance of individual sound sources, we computed power-spectral density (PSD) values separately for each participant, condition (*correct* versus *incorrect*) and the encoding and maintenance stage. To maximize reproducibility of the evoked response across trials, we applied DSS (de Cheveigné and Simon, 2008; de Cheveigné and Parra, 2014). For each participant, the first two DSS components (i.e., the two ‘most reproducible’ components; determined from the encoding stage, i.e. from 0 to 2500 ms relative to scene onset) were retained and used to project the data from the entire trial back into sensor-space. Similar analyses in which only the first, the first three, or the first twelve components were retained, yielded the same pattern of results and are not reported here. We additionally analysed the data from the maintenance stage without prior application of DSS, since it can be argued that entrained activity might show increased temporal variability across trials and participants after stimulus presentation has ended. Thus, only retaining evoked components might be detrimental to observing entrainment effects during the maintenance stage. However, the results were highly similar to those following DSS and are thus not reported.

To increase the spectral resolution of the PSD analysis, we next concatenated all trials into a single timeseries, separately for each participant, condition, and stage. This procedure results in a maximum frequency resolution of 0.02 Hz. PSD values were computed from 1 to 21 Hz by applying a Fast Fourier Transform with a single Hanning taper. The power at each frequency bin was normalized by dividing it by the mean power across the 3rd to 20th neighbouring bin above and below. For each participant and frequency, we then selected the subset of 10 channels which showed the largest normalized PSD across both the *correct* and *incorrect* conditions. Since no stimulus-related PSD peaks were observed in the maintenance stage, subsequent analyses focused on the encoding interval. For statistical comparison, we calculated an ANOVA using the variables Condition (*correct* versus *incorrect*) and rate (3, 7, 19 Hz). Finally, we estimated the sources underlying the phase locked responses at the stimulation frequencies of 3, 7 and 19 Hz, using the minimum-norm method. Data for the encoding stage (from 500 to 2500 ms, i.e. excluding the evoked response from stimulus onset) were inverted independently for 3, 7, and 19 Hz, but pooled across *correct* and *incorrect* trials. After summarizing source power in each time-frequency range of interest for each participant using a Morlet wavelet projector, source statistics maps were computed by testing the activity in each frequency band against 0.

## 3. Results

### Behavioural responses suggest that search of STM is affected by confidence

Figure 3 shows response times (RTs) and performance for each behavioural outcome, both pooled across and separately for the three probe rates (Figure 3A and B, resp.).

Statistical analysis was performed via a 2-way repeated measures ANOVA using the variables outcome (TP, TN, FP, FN) and rate (3, 7, 19 Hz).

For performance (Figure 3A and B, left), we observed a main effect of outcome (F(3, 66) = 89.65, p < .001), due to participants producing more *correct* (TP and TN) than *incorrect* (FP, FN) responses.

For RTs, we observed a main effect of outcome (F_3, 66_ = 20.26, p < .001). As evident from Figure 3A, participants were slower in TN compared to TP trials (t_22_ = -6.18, p < .001), consistent with the notion that TN responses are likely to be preceded by an exhaustive search of ASTM contents causing longer RTs, whereas TP responses are not. Participants were also slower in FP compared to TP trials (t_22_ = -10.36, p < .001; despite responding ‘same’ in both cases) and faster in TN compared FN trials (t_22_ = −2.34, p = .029.; despite responding ‘different’ in both cases). This suggests that some aspect of the cognitive processes underlying the ASTM comparison (e.g. the STM representation, the sensory representation of the probe stimulus, the decision process) was prolonged in *incorrect* trials, on average leading to delayed responses. This is also supported by the different distribution of RTs for TP and TN compared to FP and FN trials in the probability density function in Figure 3C).

In summary, the behavioural results indicate that the present task is well suited to tap into auditory short-term memory. Participants performed above chance and gave significantly more *correct* (i.e. TP and TN) than *incorrect* (i.e. FP and FN) responses in all conditions. Further, the fact that RTs were faster for *correct* compared to *incorrect* responses already points towards a potentially facilitated STM comparison- and decision-making process during *correct* trials.

### Behavioural responses reveal perceptual dominance of the 3 Hz rate

Examining performance whilst including rate as a factor (Figure 3B) we further observed a significant interaction between the variables *outcome* and *rate* (F_6, 132_ = 41.07, p < .001). This interaction was driven by the 3 Hz condition (Figure 3B left), with follow-up t-tests demonstrating more TP and FP responses for 3 Hz compared to 7 Hz (t_22_ = 10.32, p < .001; t_22_ = 5.78, p < .001) and 19 Hz trials (t_22_ = 8.41, p < .001; t_22_ = 5.17, p < .001), and fewer TN and FN responses for 3 Hz compared to 7 Hz (t_22_ = -5.78, p < .001; t_22_ = -10.32, p < .001) and 19 Hz trials (t_22_ = -5.17, p < .001; t_22_ = -8.41, p < .001). This suggests a tendency to respond with ‘same’ in trials, in which a 3 Hz probe was presented, in both ‘same’ and ‘different’ trials (see Figure 2).

For reaction times (Figure 3B) an ANOVA similarly revealed a significant interaction for RTs between the variables outcome and rate (F_6, 132_ = 7.36, p < .001). Looking at Figure 3B, this interaction was clearly driven by the 3 Hz probe condition. Follow-up t-tests confirmed that RTs to 3 Hz probes were significantly faster than those to 7 Hz and 19 Hz probes for TP (t_22_ = -7.73, p < .001; t_22_ = -6.67, p < .001) and FP outcomes (t_22_ = 3.20, p =.004; t_22_ = 3.78, p = .001), and slower than those to 7 Hz and 19 Hz probes for TN (t_22_ = -2.2, p = .039; t_22_ = -2.45, p = .023) but not for FN outcomes (p’s > .20). In other words, 3 Hz probes led to faster RTs whenever participants deemed the probe the same as the memorized stimulus, but to slower RTs when they correctly identified it as different.

Overall, this pattern of results suggests that the 3 Hz rate is perceptually dominant and likely un-bound from its carrier frequency such that participants are biased to respond that the probe was present even if it is presented with the incorrect carrier frequency (i.e it is a “different” trial).

### MEG data reveal sustained increases in power during *correct* compared to *incorrect* trials

Figure 4 displays the group-level Event related field (ERF) results. The presentation of the ‘scene’ stimuli evoked increased sustained activity that persisted for the duration of the encoding stage. This was followed by a drop in sustained power, during the maintenance stage. Activity associated with the onset of the probe, and subsequent button press, is seen during the retrieval stage. In the following we focus on understanding how activity during the Encoding and Maintenance stages (i.e. before the probe has been presented) differs between *correct* and *incorrect* trials.

We observed patches of significantly larger instantaneous power (RMS across all channels) for *correct* compared to *incorrect* trials, as determined via a bootstrapping procedure, throughout most of the encoding and maintenance stages (Figure 4A). This difference is also evident in the fieldmaps (Figure 4B), whose dipolar topography suggests an underlying auditory source. A cluster-based permutation analysis identified a left-hemispheric channel cluster, during the encoding-(p = .037; 21 channels) and a left-hemispheric and centrally located cluster during the maintenance stage (p = .011; 33 channels), indicated in Figure 4B. Consistent with the RMS results, event-related field data from those channels (Figure 4B) show higher activity during trials which were ultimately *correct*. Strikingly, increased activity for correct trials persists in the maintenance stage, even when sound is no longer presented.

### Differences between *correct* and *incorrect* trials are restricted to auditory areas during the encoding stage, shifting to non-auditory areas during the maintenance stage

Source inversion based on multiple-sparse priors (Figure 4C) suggest the above observed differences arise predominantly in bilateral temporal areas during the encoding stage, in line with stronger auditory cortex activation during *correct* compared to *incorrect* trials. During the maintenance stage, we found bilateral differences in parietal areas as well as prefrontal and temporal areas. Together, these results indicate that neural activity differentiating *correct* from *incorrect* trials during the encoding stage is predominantly restricted to auditory sensory areas. Subsequently, during short term maintenance, this shifts to other, non-auditory areas.

### No differences in Power-spectral density between *correct* and *incorrect* trials

Figure 5 illustrates the results of the PSD analysis. During the encoding stage (Figure 5A,), distinct peaks are visible at the stimulation frequencies of 3, 7, and 19 Hz and their harmonics. However, no representation of the stimulus frequencies was visible during the maintenance stage (not shown in Figure 5). Focusing on the encoding stage, Figure 5B displays the peak PSD values for each rate and outcome conditions. The ANOVA using the variables outcome (*correct* versus *incorrect*), and AM rate (3, 7, 19 Hz) yielded a significant main effect of AM rate (F_2, 44_ = 12.35, p < .001). Previous work (Elhilali et al., 2009; Huang and Elhilali, 2020) has revealed large scale effects of attention on PSD measures of auditory tracking such that attended sounds are associated with increased sustained (steady-state) neural representation. The absence of a difference between *correct* and *incorrect* trials here suggests that success or failure in the present task is not driven by variable attentive tracking of the scene components. Source level estimation of the PSD at 3 Hz (tested against 0; all displayed voxels are p < 0.05) shows extensive activation with a bilateral peak in temporal regions, in line with auditory sensory activation (Figure 5C). Source estimations of the 7 and 19 Hz components yielded highly similar statistical maps (not shown).

**Figure 5.**
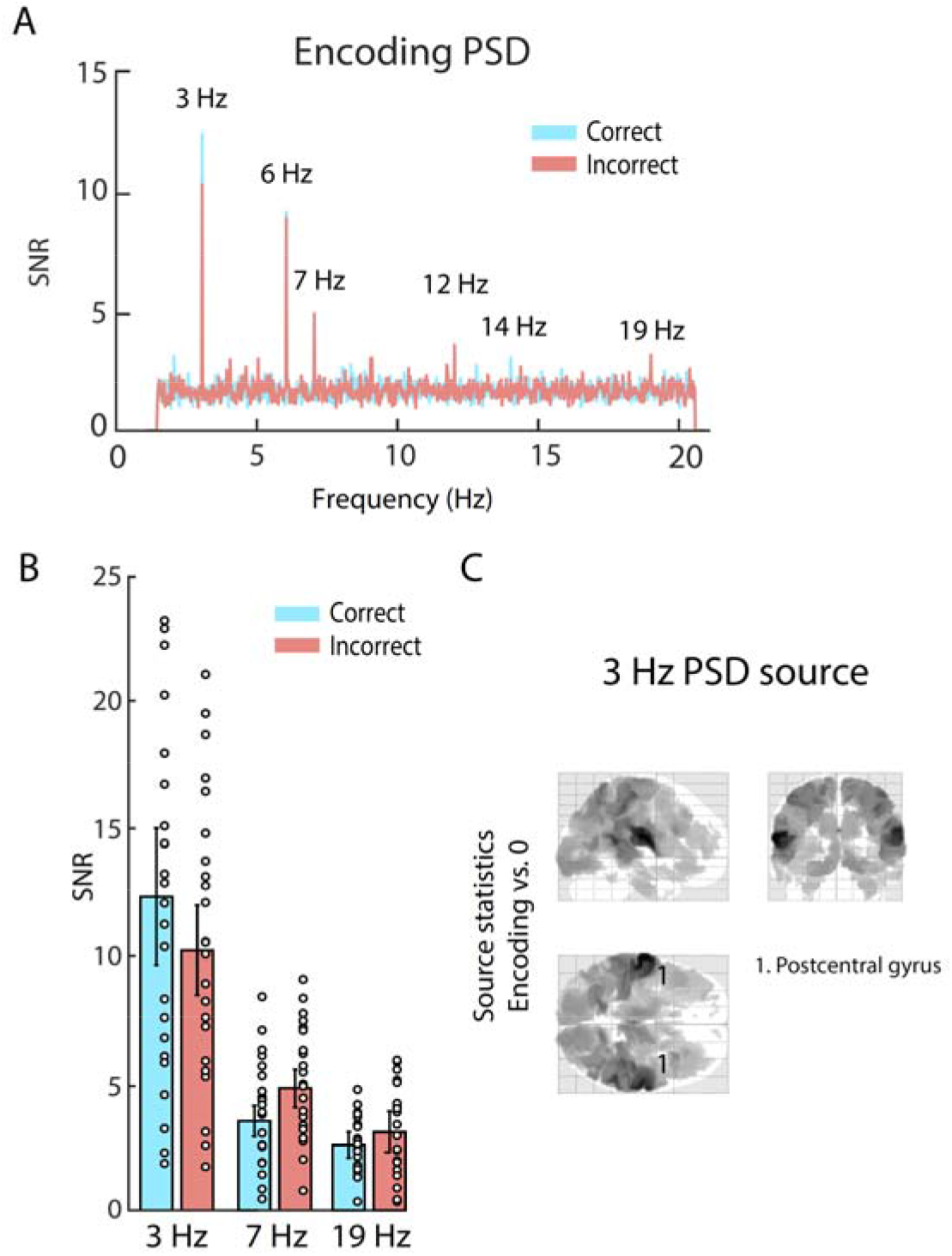
Power-spectral density (PSD). **(A)** Normalized PSD during the encoding phase (no noticeable peaks were found during the maintenance stage). Blue represents data from *correct* trials, red data from *incorrect* trials. Numbers highlight peaks at stimulation frequencies as well as harmonics. **(B)** PSD values for the three rates (3 Hz, 7 Hz, 19 Hz) for *correct* (blue) and *incorrect* (red) trials. Circles indicate individual participants, error bars represent SEM. **(C)** Source-level difference in oscillatory power at 3Hz, between *correct* and *incorrect* trials during the encoding time window. The number indicates the significant cluster (p<0.05), the inset indicates the anatomical area containing the cluster’s peak voxel.

### Widespread increases in oscillatory alpha, beta, and theta power during *correct* trials

Next, we looked into differences between *correct* and *incorrect* trials, separately within the alpha, beta and theta frequency bands. Figure 6 displays time-frequency planes and source inversions of the observed effects in the alpha-band. At scalp level (Figure 6A), distinct increases in power relative to the baseline can be seen, which appear to be stronger in the *correct* compared to the *incorrect* condition. Using cluster-corrected permutation tests in time and space, we observed significantly larger power for *correct* versus *incorrect* trials, during both the encoding (p = .014, 1500 – 2500 ms, 137 channels) and maintenance stages (p = .004, 2500 – 3650 ms, 117 channels). Source inversion of the respective time-frequency windows (Figure 6B) revealed increased activity for *correct* versus *incorrect* trials in left parietal areas during the encoding stage, and in a more widespread parieto-occipital area as well as a frontal region during the maintenance stage.

**Figure 6.**
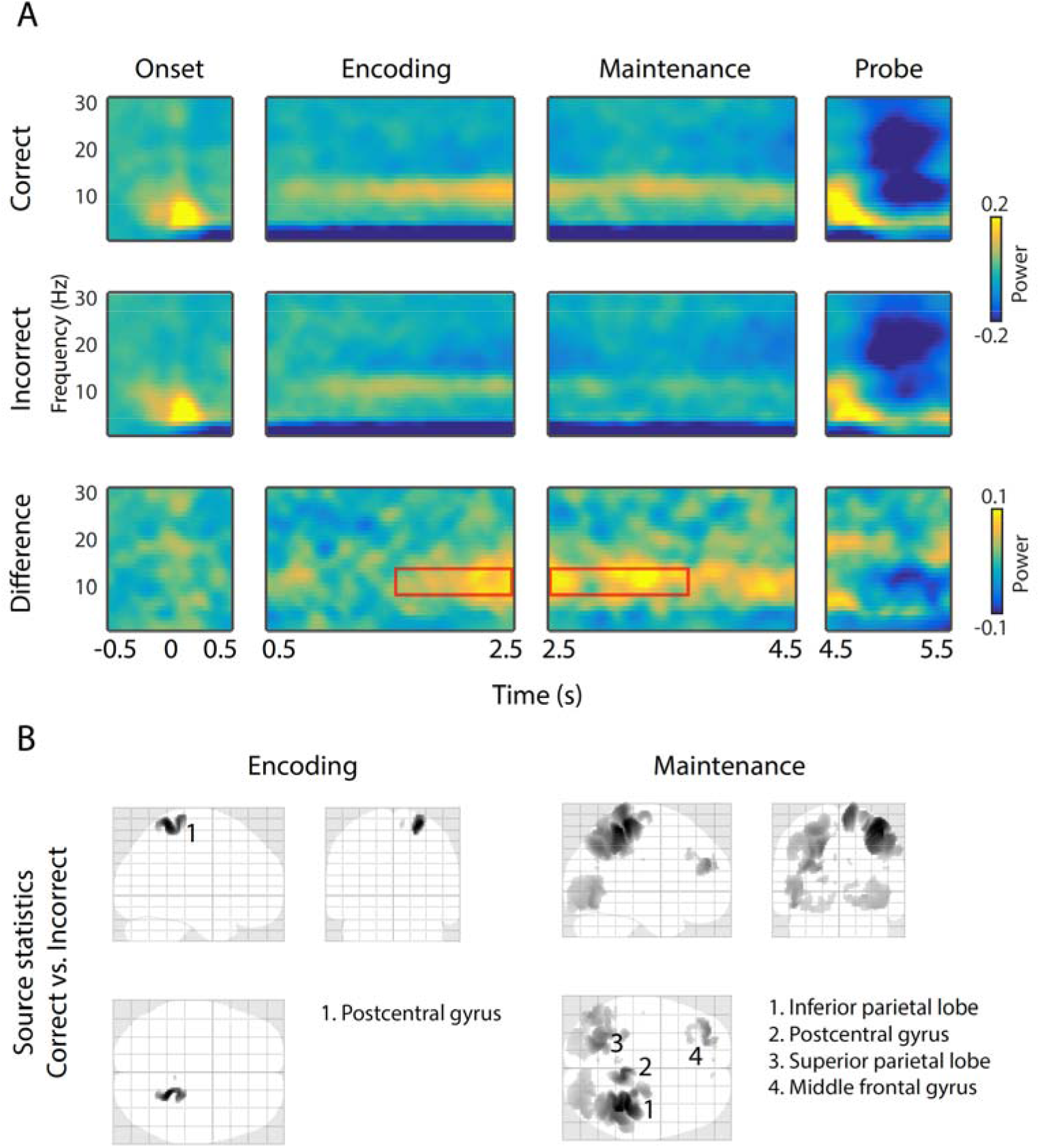
Results of alpha-band (8-14 Hz) time-frequency analysis. **(A)** Time-frequency representations of oscillatory power during the stimulus onset, encoding, maintenance, and probe phase of the experiment. Data are shown separately for *correct* (top) and *incorrect* (middle) trials, as well as their difference (bottom). Red boxes indicate time-windows exhibiting significant differences, as tested via a cluster-based permutation procedure. The plotted data represent the mean across all sensors belonging to the significant cluster. **(B)** Source-level difference between *correct* and *incorrect* trials during the time windows determined via scalp analysis, separately for the encoding (left) and maintenance stage (right). Numbers indicate significant clusters (p<0.05), sorted by p-values from low to high. Insets indicate anatomical areas containing the peak voxel of each cluster.

Figures 7 shows time-frequency results and source inversions of the observed effects in the theta- and beta-bands. In the theta-band, we found significantly higher power in *correct* compared to *incorrect* trials, during the second half of the encoding stage (p = .031, 1560 – 2410 ms, 93 channels). Source inversion of this time-frequency window yielded increased activity in left temporal regions for the *correct* versus the *incorrect* condition.

**Figure 7.**
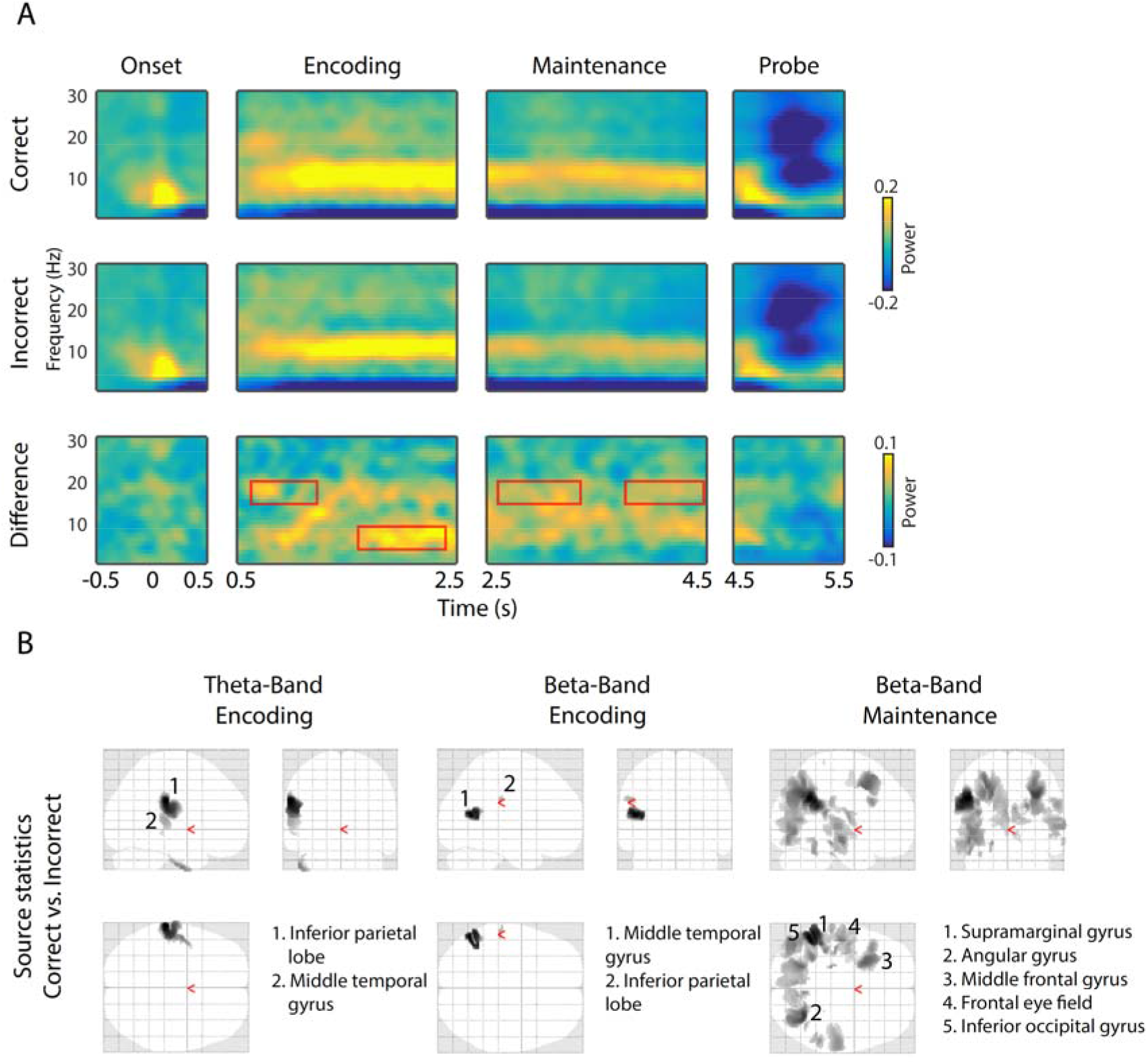
Results of theta (4-7 Hz) and beta-band (15-20 Hz) time-frequency analysis. **(A)** Time-frequency representations of oscillatory power during the stimulus onset, encoding, maintenance, and probe phase of the experiment. Data are shown separately for *correct* (top) and *incorrect* (middle) trials, as well as their difference (bottom). Red boxes indicate time-windows exhibiting significant differences, as tested via a cluster-based permutation procedure. The plotted data represent the mean across all sensors belonging to the significant cluster. **(B)** Source-level difference between *correct* and *incorrect* trials during the time windows determined via scalp analysis, separately for the theta-band encoding stage (left), and the beta-band encoding (middle) and maintenance stages (right). Numbers indicate significant clusters (p<0.05), sorted by p-values from low to high. Insets indicate anatomical areas containing the peak voxel of each cluster.

In the beta-band, we again found higher power in *correct* versus *incorrect* trials, however during an earlier time-window (p = .02, 646 – 1281 ms, 80 channels) in the encoding stage, as well as during two extended time-windows in the maintenance stage (p = .005, 2632 – 3340 ms, 152 channels and p = .007, 3766 – 4510 ms, 140 channels). For the effect during the encoding time-window, source inversion revealed a likely source in left temporal areas, similar to the theta-band. For the source inversion of the effect during maintenance, we inverted a single time stage spanning the time-windows of both clusters. Here, we found increased activity for *correct* versus *incorrect* trials in widespread left-temporal and occipital areas, overlapping with the areas observed for the other theta- and beta-band effects.

Taken together, oscillatory power in both the alpha and beta bands revealed more widespread differences during maintenance compared to encoding, yet in different cortical areas. Interestingly, these differences were largely outside of auditory sensory areas. This is in contrast to the effects observed in the encoding time window for beta and theta bands, whose estimated sources were in or close to left-hemispheric auditory areas. Given the fact that the time-windows and/ or estimated source of the significant effects were different between the three investigated frequency bands, it is unlikely that they are due to a broad-band increase in power during *correct* trials. Rather, we suggest that each frequency band is part of a distinct functional mechanism, which jointly contribute to successful task performance.

## 4. Discussion

Auditory short-term memory plays a critical role in our daily life, from communication, to scene analysis and music appreciation. Understanding the brain mechanisms that support auditory memory is important for elucidating fundamental brain functions and understanding individual differences and impairments, e.g. such as those arising in the course of aging (Bopp & Verhaeghen, 2005; Cooper et al., 2006; Rimmele et al., 2012; Ruzzoli et al., 2012; Sur & Golob, 2020). Previous work into auditory short-term memory predominantly focused on contrasting tasks of different difficulty (i.e. high vs low memory load). Here we ask a somewhat different question – what are the neural correlates that distinguish ultimately successful versus unsuccessful short-term retention of a complex auditory scene?

We employed a novel ASTM task (Figure 2), which required listeners to encode and retain three simultaneously presented artificial auditory objects, defined by frequency and rate. Thus, the proper execution of the STM task necessitates the *correct* and sustained binding of the frequency and rate of each of the three auditory streams, and failure to do so potentially underlies *incorrect* task performance (Bizley and Cohen, 2013; Darwin, 1997; Joseph et al., 2016, 2015). We analysed activity during the encoding and maintenance stages of the task (i.e. before the probe stimulus was presented) to determine what patterns of activation might be associated with ultimately *correct* or *incorrect* responses to the probe. Our results reveal a distributed pattern of enhanced ERFs and oscillatory neural activity in several frequency bands specific to encoding and maintenance stages, and indicative of successful short-term memory performance.

### STM search and confidence effects revealed in behavioural response patterns

Overall, listeners demonstrated an average hit-rate of 71%, confirming a challenging behavioural task. We observed faster RTs for TP (True positive; i.e when the probe was present in the scene and identified as such) versus TN responses (True negative; when the probe was absent from the previously heard scene and identified as such), which suggests a non-exhaustive scan of STM storage in cases participants correctly determined the probe to be present in the just heard scene. Participants were also faster during *correct* compared to *incorrect* trials, indicative of higher levels of response confidence in the former (Kahana and Loftus, 1999; Murdock, 1985; Weidemann and Kahana, 2016).

### Behavioural responses suggest perceptual salience of the 3 Hz rate

Interestingly, performance for trials containing 3 Hz probes stood out in two aspects. Firstly, mean RTs were faster for 3 Hz compared to 7 Hz and 9 Hz probe trials when participants gave a *same* response (i.e. TP, FP), regardless of whether the outcome was *correct* or *incorrect*. The opposite was true for TN trials, in which they gave a *different* response. Secondly, participants gave significantly more *same* than *different* responses for 3 Hz compared to 7 or 19 Hz probe trials. Again, this was the case regardless of whether the outcome was *correct* or *incorrect* and despite the fact that participants received training and feedback after every trial. This performance pattern indicates that slow repetition rates (3 Hz here) might be perceptually more salient than faster rates. That is, they might pop out perceptually and attract attention, but, as discussed below, depending on the task context might ultimately impede performance.

Previous works have highlighted differences in the natural occurrence of, and processing mechanisms for auditory stimuli with slow compared to fast amplitude modulation (e.g. Poeppel, 2003; Giraud and Poeppel, 2012). One hypothesis is that this reflects the tuning of the auditory system to the syllabic rate of spoken language (∼4 Hz; Chandrasekaran et al., 2009; Chi et al., 1999; Elliott and Theunissen, 2009). Notably, here 3 Hz probes were also more often incorrectly identified as being present in the scene. This suggests that rate and frequency information were not strictly bound in these sounds, and that the 3 Hz rate information dominated the perceptual representation even though the task required participants to remember the rate in association with the carrier. These incidental observations speak to a longstanding question (e.g. see reviewed in Sohoglu et al, 2020) regarding whether spectral and temporal acoustic features are represented independently, or in an integrated manner.

### No differences in Power-spectral density between *correct* and *incorrect* trials during the encoding period

The analysis of brain responses focused on the encoding and maintenance stages – i.e before the presentation of the probe (and the ensuing behavioural response). This allowed us to pinpoint the aspects of brain activity that were most different between subsequently successful or failed trials.

As expected, we found spectral peaks at the three AM stimulation frequencies (3, 7, 19 Hz and harmonics) during the encoding stage. In line with the observed ERFs (see below), these were localized in or close to primary auditory cortex. The spectral amplitude at 3 Hz was significantly larger compared to 7 and 19 Hz. This matches the behavioural findings of overall faster responses as well as more *same* than *different* responses for trials with 3 Hz probes and suggests an increased saliency of 3 Hz compared to 7 and 19 Hz stimuli.

Contrary to our expectations however, PSD did not differ between subsequently *correct* and *incorrect* trials. In the visual domain, such a differentiating neural pattern has been observed during the STM maintenance of several stimuli (Killebrew et al., 2018; Peterson et al., 2014), and one study found a comparable result using only a single auditory stimulus stream (Falk et al., 2017). As stated above, and given the many demonstrations that focused attention is associated with increased SNR at the stimulus rate (Ding and Simon, 2013; Elhilali et al., 2009; Mesgarani and Chang, 2012; Riecke et al., 2014; Xiang et al., 2010; Zion Golumbic et al., 2013) the absence of a difference between *correct* and *incorrect* trails here suggests that stimulus-directed attention per se is not what affected success or failure in the present task. Namely, *correct* trials are not simply those trials during which participants tracked the unfolding auditory streams more intently during the encoding stage.

Further, we observed no persistence of stimulus rate representation into the maintenance stage. In principle, such a continuation of rhythmic neural activity beyond the offset of the entraining auditory stimulus is possible, as two recent studies have demonstrated (Kösem et al., 2018; van Bree et al., 2020). A likely reason for the current lack of observable PSD effects lies in the more complex stimulus characteristics (i.e. a mix of three simultaneous sound streams instead of a single stream used in previous work) and consequent potential variability in rehearsal strategies across trials and participants.

### *Correct* trials are associated with increased event-related activity relative to *incorrect* trials

Starting early on during the encoding stage and continuing throughout the maintenance stage, we observed increased ERF activity for *correct* compared to *incorrect* trials, particularly at temporo-parietal sensors. Previous studies have reported analogous increases in event-related EEG and MEG activity during the maintenance period of high compared to low load ASTM tasks (Grimault et al., 2014; Huang et al., 2016; Nolden et al., 2013). Similarly, Cano and Knight (2016) observed stronger ERP responses in a think/ no think paradigm, where participants tried to remember an auditorily presented word pair compared to when they tried to suppress it. In a cross-species comparison between humans and monkeys, Huang et al. (2016) provided evidence that such increased ERF activity likely originates from elevated firing of neurons in auditory cortex. This is in line with our observed source activity particularly during the encoding stage, in which *correct* compared to *incorrect* trials showed stronger activity in or very close to primary auditory cortex (see also Albouy et al, 2013; Kumar et al., 2016, 2021).

During maintenance, source level activity showed a more widely distributed pattern. In particular, we observed stronger ERFs for *correct* compared to *incorrect* trials in the dorsolateral prefrontal cortex (DLPFC), a key structure in visual and auditory short-term memory operations (Barbey et al., 2013; Bodner et al., 1996; Goldman-Rakic, 1995) and sustained attentional control (Kane and Engle, 2002). Load-related increases in DLPFC activity were previously found in an MEG study during a non-verbal auditory STM task (Grimault et al., 2014) as well as in a fMRI study specifically dissociating attention- and load related activations during auditory STM (Huang et al., 2013).

Rather than the actual storage of information, recent models have proposed that PFC activity during short-term memory operations likely reflects executive top-down signals to sensory areas concerned with maintenance (Lara and Wallis, 2015; Postle, 2006; Sreenivasan et al., 2014) as well as rehearsal strategies (Awh et al., 1996; D’Esposito et al., 2000). Similarly, our further observed parietal ERF sources (superior parietal lobe, postcentral gyrus) have previously been associated with motor sequencing during speech preparation and/ or production (Dhanjal et al., 2008; Heim et al., 2012). Although the present stimuli are unlikely maintained in the same way as speech, this hints at a potential involvement of silent rehearsal processes, especially during successful maintenance (see also Nolden et al., 2013).

Together, the ERF results are consistent with previous demonstrations of persistent activity during encoding and maintenance of auditory short-term memories (Fuster and Jervey, 1982; Huang et al., 2016; Constantinidis et al., 2018; but see also Masse et al., 2020). Critically, unlike the previous investigations reviewed above where the stimuli/task load were manipulated to affect ASTM performance, here task difficulty was kept fixed throughout the experiment. The activations observed suggest that parietal activity is also modulated on a trial by trial basis. This activity likely either directly represents the memorized information or pertains to supporting mechanisms for rehearsal or suppression of distractors during the maintenance period that ultimately result in a successful vs unsuccessful performance.

### Successful ASTM performance relies on distinct sources of alpha-band activity in posterior and fronto-parietal regions

As hypothesized, we observed increased ongoing alpha-band activity in *correct* compared to *incorrect* trials, during both encoding and maintenance stages. Particularly the latter exhibited power differences in a large area around right posterior parietal regions. This extends previous studies, which reported general increases in posterior alpha-band activity during ASTM operations compared to rest (Kawasaki et al., 2010; Luo et al., 2005), as well as parametrically increasing with STM load (Obleser et al., 2012; Wisniewski et al., 2017). For example, Obleser et al. (2012) had participants memorize different numbers of spoken digits and found that alpha-band power during the maintenance interval increases with memory load. This memory-related increase in posterior alpha-power has also consistently been shown for other sensory modalities (Jensen et al., 2002; Spitzer and Blankenburg, 2012; Van Dijk et al., 2010) and is widely suggested to reflect functional inhibition of task irrelevant cortical areas (Jensen and Mazaheri, 2010; Klimesch et al., 2007; Wilsch and Obleser, 2016). That is, alpha-band mediated inhibition ensures that sufficient processing resources are allocated for attending to and maintaining auditory stimulus information. Our data further develops this line of research by demonstrating, that a certain level of functional inhibition via alpha-band power increase seems necessary for successful ASTM performance. Since no additional salient external sensory inputs were present in our experiment, the observed posterior alpha activity likely reflects protection from internal rather than external noise.

Interestingly, in addition to the posterior increases of alpha power associated with functional inhibition, our data also show a frontal as well as a temporo-parietal source. The latter, which comprised the strongest source of alpha-band power differences in our data, has repeatedly been found to be activated during ASTM (Gaab et al., 2003; Obleser et al., 2012; Paulesu et al., 1993; Wilsch et al., 2015). For instance, in an fMRI study, Gaab et al. (2003) observed increased activity in the supramarginal gyrus, overlapping with our temporo-parietal source, during a pitch retention task, which correlated with task performance. More generally, an increasing number of studies have reported corresponding increases in alpha activity outside of sensory areas during STM operations (Leiberg et al., 2006; Palva et al., 2010; Wilsch et al., 2018). These activations hint at an important role of alpha in more executive functions and reflecting aspects of top-down modulation of sensory areas during short-term memory operation, in order to temporally maintain sensory representations (Wilsch and Obleser, 2016). In this light, our results extend existing knowledge by demonstrating that successful ASTM operations rely, at least partly, on two distinct types of alpha-band activity: One originating in posterior regions, concerned with functional inhibition, and one originating in fronto-parietal regions, involved in executive control of sensory maintenance.

### Increased left parietal theta-band activity during the encoding stage differentiates *correct* from *incorrect* trials

In contrast to the alpha- and beta-band, we observed effects in theta-band activity only during the encoding stage where activity was enhanced during subsequently *correct* compared to *incorrect* trials. These effects arose partway through the trial and therefore may be related to fatigue or random failure of the underlying STM encoding mechanisms during the *incorrect* trials.

In visual STM tasks, theta activity has been prominently observed in a large number of studies, both in posterior sensory- and association areas (Costers et al., 2020; Raghavachari et al., 2006, 2001) as well as in frontal-midline regions (Jensen and Tesche, 2002; Kawasaki et al., 2010; Roux and Uhlhaas, 2014; Ede et al., 2017; Meltzer et al., 2017; Proskovec et al., 2018). While there is less known regarding the role of theta in STM outside the visual domain (but see e.g. Cano and Knight, 2016), recent studies have demonstrated an increase in ASTM performance by enhancing ongoing cortical theta activity during maintenance using TMS (Albouy et al., 2017) and rhythmic visual stimulation (Albouy et al., 2022). Based on an a-priori measurement of STM related neural activity via EEG and MEG, their site of stimulation was located over the left intraparietal sulcus. Notably, our presently observed source for the contrast of theta power between *correct* and *incorrect* trials was in the same region, around the left supramarginal gyrus, as well as the left middle temporal gyrus.

Unlike the present results, the theta-band effect from Albouy et al. (2017) was present in the maintenance stage. While likely tapping into the same neural mechanisms, a possible reason for this difference is that the task used by Albouy et al. (2017) required active manipulation of STM contents compared to a simple maintenance in the present study. Using fMRI, other studies have also found evidence for auditory STM related operations in this region, especially during highly demanding tasks (Foster et al., 2013; Foster and Zatorre, 2010; Zatorre et al., 2010). Additionally, the present effects in the alpha and beta band power were also located around the supramarginal gyrus, further supporting its importance in auditory STM operations.

### Increased beta-band power during maintenance may relate to binding of ASTM contents in sensory areas

Corresponding to our results in the alpha- and theta band, beta-band power was elevated during *correct* versus *incorrect* trials.

Compared to other frequency bands, the role of beta-band activity during STM, and in particular for auditory information, has been considerably less studied. In line with our data, Leiberg et al. (2006) found that beta-band activity in temporal areas increases parametrically with auditory STM load. In addition, several previous studies have reported analogously increased levels of beta-band activity during visual STM tasks, predominantly in visual sensory areas (Chen and Huang, 2016; Daume et al., 2017; Deiber et al., 2007; Fodor et al., 2018; Killebrew et al., 2018). For instance, in a recent visual short-term memory study, Fodor et al. (2018) observed reduced beta-band activity in participants with mild cognitive impairment compared to healthy controls, along with reduced STM performance. Further, linking structural MRI to the EEG measures revealed that the strength of beta-band activity correlated significantly with the size of the hippocampus, entorhinal cortex and parahippocampal cortex, all of which are suggested to be involved in STM operations (Axmacher et al., 2008; Kumar et al., 2016). The fact that we observed more pronounced and widespread differences between *correct* and *incorrect* trials during the maintenance stage might suggest, that beta-band is particularly involved in the preservation of sensory information. In line with this interpretation, Weiss and Mueller (2012) speculated that beta-band activity reflects the binding of auditory short-term memory contents in sensory areas.

Spatially, the strongest difference between *correct* and *incorrect* trials in the beta band was localized around the supramarginal gyrus (BA40). This mirrors our findings in the alpha- and theta band results and adds a further indication of this regions important role in auditory STM operations (Celsis et al., 1999; Gaab et al., 2003; Paulesu et al., 1993; Salmon et al., 1996). Further significant effects encompassed the left angular gyrus and the left middle temporal gyrus. A joint increase in these two regions during auditory processing compared to a non-auditory control condition has been found by both Goycoolea et al. (2005) and Ahmad et al. (2003) using fMRI.

Finally, two additionally observed sources constitute the secondary visual cortex and the frontal eye-field. We speculate that these reflect increased suppression of potentially distracting visual information in subsequently *correct* trials.

### Conclusions

This study goes beyond previous findings by providing a thorough account of oscillatory and evoked neural activity patterns differentiating (ultimately) successful from unsuccessful ASTM performance (see Figure 8 for a summary). Notably, we show that successful compared to unsuccessful ASTM performance was associated with a significant overall enhancement of event-related fields and oscillatory activity in the theta, alpha, and beta frequency ranges in line with previous reports of persistent neural activity changes during STM. Effects in each frequency band were linked with different underlying sources, suggesting that they represent distinct functional mechanisms which jointly enable successful memory performance. Spatially, these effects were confined to posterior sensory areas during encoding and spread to parietal and frontal areas during maintenance, presumably reflecting STM contents and top-down control processes, respectively.

**Figure 8.**
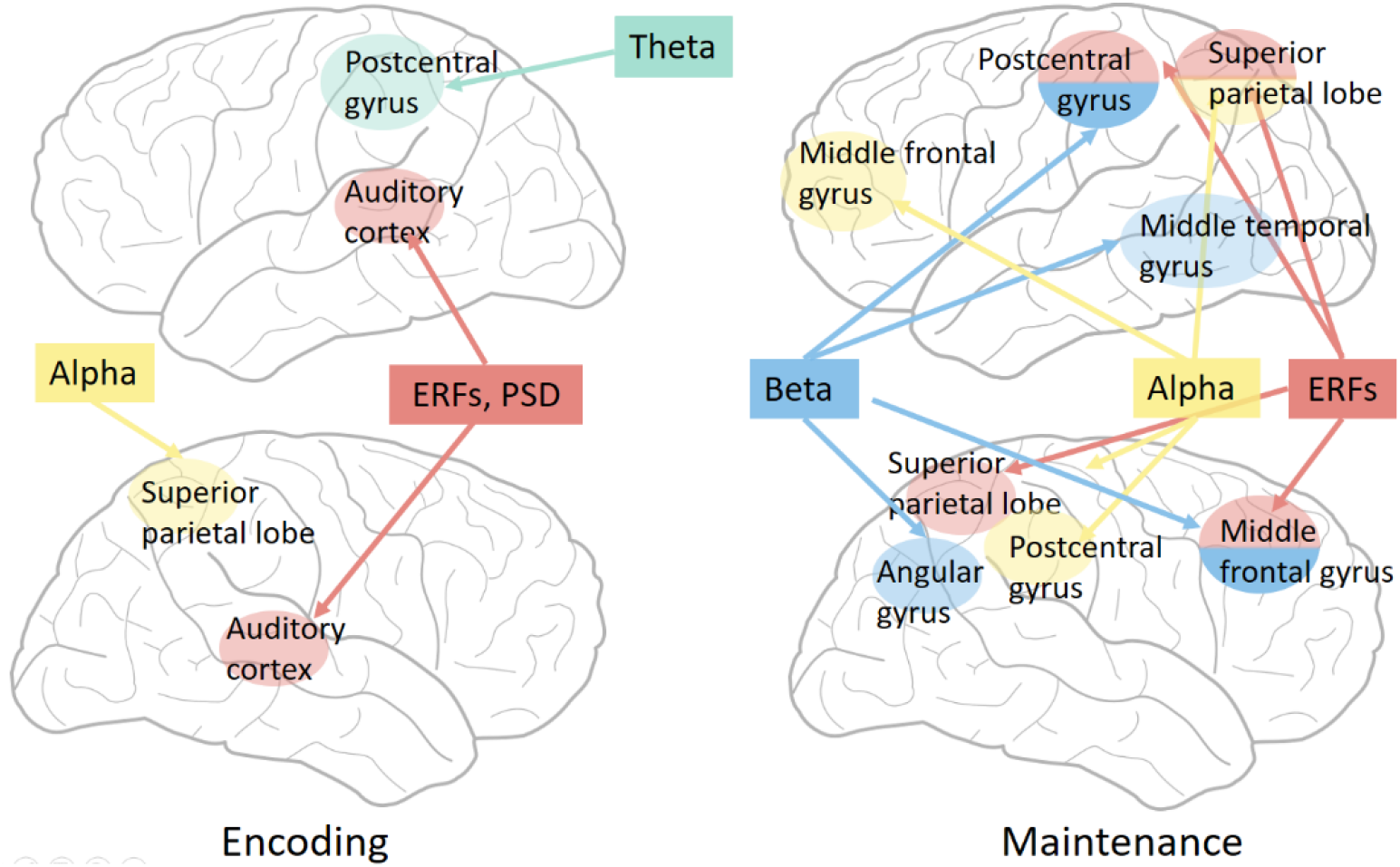
Summary of findings. Illustration of brain areas showing increased neural activity during *correct* versus *incorrect* ASTM performance, separately for encoding (left) and maintenance stages (right). ERF: Event-related fields; PSD: Power-spectral density.

## Data-availability

Data and relevant code will be uploaded to an appropriate online depository upon publication.

## Acknowledgements

This research was supported by a EC Horizon 2020 grant and a BBSRC international partnering award to MC.

